# Development of Resistance to 4’-Ethynyl-2-Fluoro-2’-Deoxyadenosine (EFdA) by WT and Nucleoside Reverse Transcriptase Inhibitor Resistant Human Immunodeficiency Virus Type 1

**DOI:** 10.1101/2020.09.04.281485

**Authors:** Maria E. Cilento, Eleftherios Michailidis, Tatiana V. Ilina, Eva Nagy, Hiroaki Mitsuya, Michael A. Parniak, Philip R. Tedbury, Stefan G. Sarafianos

## Abstract

4’-ethynyl-2-fluoro-2’-deoxyadenosine (EFdA, MK-8591, islatravir) is a nucleoside reverse transcriptase translocation inhibitor (NRTTI) with exceptional potency against WT and drug-resistant HIV strains. However, HIV resistance to EFdA is not well characterized. We therefore developed resistance to EFdA by serial passages using progressively increasing concentrations of EFdA. The starting virus was either WT or clinically relevant NRTI-resistant viruses K65R, M184V, and D67N/K70R/T215F/K219Q). In all cases, the selected mutations included M184V. Additional mutations in the RT connection domain (R358K and E399K) and one mutation in the RNase H domain (A502V) were noted. Site-specific mutagenesis validated the role for M184V as the primary determinant for resistance to EFdA; none of the connection domain mutations contributed significantly to phenotypic resistance to EFdA. A novel EFdA resistance mutation was also observed in the background of M184V. The A114S/M184V combination of mutations imparted higher resistance to EFdA (~24-fold) than M184V (−8-fold) or A114S (~2-fold) alone. Virus fitness data suggested that A114S affects HIV fitness by itself and in the presence of M184V. This is consistent with biochemical experiments that showed decreases in the enzymatic efficiency (k_cat_/K_m_) of WT RT *vs.* A114S (2.1-fold) and A114S/M184V/502V (6.5-fold), whereas there was no significant effect of A502V on RT or virus fitness. The observed EFdA resistance of M184V by itself and in combination with A114S combined with the strong published *in vitro* and *in vivo* data, confirm that EFdA is an excellent candidate as a potential HIV therapeutic.

## INTRODUCTION

As of 2018, 37.9 million people worldwide are living with HIV/AIDS, with 1.7 million new HIV infections and 770,000 AIDS-related deaths annually (Mahy et al., 2019). However, AIDS-related morbidity and mortality rates have declined in recent years, largely due to the widespread use of highly-active antiretroviral therapy (HAART) (Mahy et al., 2019). HAART typically consists of a combination of two nucleoside reverse transcriptase inhibitors (NRTIs), along with a nonnucleoside reverse transcriptase inhibitor, protease inhibitor or an integrase inhibitor (Merluzzi et al., 2010). Azidothymidine (AZT), didanosine (ddI), lamivudine (3TC), emtricitabine (FTC), abacavir (ABC), and tenofovir (TFV) are the six NRTIs included in HAART regimens (Eggleton and Nagalli, 2020; Menéndez-Arias, 2008; Merluzzi et al., 2010). However, the prevalence of HIV strains resistant to these compounds is rapidly increasing, both in treatment-experienced and newly-infected patients (Clutter et al., 2016; Eggleton and Nagalli, 2020; Larder et al., 1995; Little et al., 2002; Pennings, 2013; Wainberg et al., 2011). High-level resistance to AZT generally requires multiple mutations, including D67N, K70R, T215F, and K219Q (Kellam et al., 1992; Larder and Kemp, 1989; Menéndez-Arias, 2008; Nakata et al., 2007), while an M184I/V mutation grants resistance to both 3TC and FTC (Menéndez-Arias, 2008; Petrella et al., 2004; Schinazi et al., 1993; Tisdale et al., 1993). Meanwhile, the K65R mutation imparts some resistance to TFV (Brenner and Coutsinos, 2009; Margot et al., 2002; Miller, 2004; Naeger and Struble, 2006). Also, of great concern is the emergence of virus strains with cross-resistance to multiple NRTIs, which can limit treatment options following viral escape from first-line HAART (Eggleton and Nagalli, 2020). M184V strains show resistance to ddI and abacavir, while mutations selected by AZT are resistant to FTC, and K65R has reduced sensitivity to ddI, 3TC, and FTC (Bazmi et al., 2000; Brenner and Coutsinos, 2009; Menéndez-Arias, 2008; Miller, 2004; Zhang et al., 1994). The currently available NRTIs can also display toxicity and side effects (Brinkman et al., 1999, 1998; Brinkman and Kakuda, 2000; Eggleton and Nagalli, 2020; Lewis et al., 2003). Additionally, the success of antiretroviral regimens at preventing HIV-1 infection has moved attention to the question of compliance, and increased the interest in therapeutic agents that may be suitable for long interval dosing (Cihlar and Fordyce, 2016; Margolis and Boffito, 2015). All of the above are reasons there is a need for the development of novel therapeutic agents.

Several studies have investigated a group of novel 4’-substituted NRTIs, the most promising of which is 4’-ethynyl-2-fluoro-2’-deoxyadenosine (EFdA, MK-8591, or islatravir) (reviewed at (Markowitz and Sarafianos, 2018)). EFdA is a deoxyadenosine analog with an ethynyl group at the 4’ carbon of the ribose and a fluorine on the 2-position of the adenine base (Kageyama et al., 2011; Kirby et al., 2013, 2011; Nakata et al., 2007). EFdA also retains a 3’-OH, unlike all other HIV NRTIs currently approved for therapeutic use. The presence of the 3’-OH improves the recognition of EFdA as a substrate by cellular kinases such as deoxycytidine kinase (Gallois-Montbrun et al., 2002; Kawamoto et al., 2008a; Nakata et al., 2007) and HIV reverse transcriptase (RT) dNTP binding domain (Michailidis et al., 2014a, 2009; Salie et al., 2016); this may contribute to the efficient production of EFdA-triphosphate (EFdA-TP) and the efficient incorporation of EFdA-monophosphate (EFdA-MP) into nascent HIV DNA during reverse transcription. Deamination of EFdA by cellular enzymes is greatly reduced by the 2-fluorine substitution, increasing the intracellular half-life of the molecule compared to similar non-halogenated compounds (Kawamoto et al., 2008a; Kirby et al., 2013). The 4’-ethynyl group of EFdA interacts with a hydrophobic pocket in the active site of RT, affecting translocation and extension of the DNA primer, despite the available 3’-OH (Michailidis et al., 2014b; Muftuoglu et al., 2014; Salie et al., 2016). Therefore, EFdA is termed a nucleoside RT translocation inhibitor (NRTTI). EFdA has demonstrated a high level of potency against both WT (wild-type) HIV-1, HIV-2, and NRTI resistant strains (Kawamoto et al., 2008a; Maeda et al., 2014; Oliveira et al., 2017; Wu et al., 2017). The NRTI-resistant virus, K65R, showed hypersensitivity to EFdA (Maeda et al., 2014; Michailidis et al., 2013). EFdA has shown *in vivo* potency against both HIV and SIV in humanized mouse (Hattori et al., 2009; Shanmugasundaram et al., 2016; Stoddart et al., 2015) and non-human primate models, respectively (Markowitz et al., 2020; Murphey-Corb et al., 2012; Stoddart et al., 2015). EFdA imparts minimal toxicity in the animal models as well as in all *in vitro* assays, due to minimal inhibition of human DNA polymerases, leading to a high selectivity index (Shanmugasundaram et al., 2016; Sohl et al., 2012; Stoddart et al., 2015). Perhaps most importantly, EFdA is being tested for once-week and once-monthly dosing regimens (Barrett et al., 2018; Grobler et al., 2019; J. et al., 2017, 2016). Taken together, these results suggest that EFdA has great promise as a potential therapeutic agent.

Little is known about the capacity of HIV to develop high-level resistance to EFdA, and what mutations may allow the virus to escape inhibition. M184V in RT decreases sensitivity to EFdA (Kawamoto et al., 2008a; Kodama et al., 2001; Maeda et al., 2014; Nakata et al., 2007; Oliveira et al., 2017; Yang et al., 2008). M184V, along with I142V and T165R, was also selected during passage of WT virus with the parental compound, EdA (Kawamoto et al., 2008a). While an I142V/T165R/M184V virus had a 22-fold increase in resistance to EFdA relative to WT, it is uncertain whether these mutations would arise during passage of virus in EFdA itself, or if novel mutations conferring even greater resistance are possible. It is also not well understood how resistance to EFdA develops in virus strains with specific pre-existing NRTI resistance mutations, which would be relevant to its potential value as salvage therapy for patients failing first-line HAART. Finally, there is no significant information on the biochemical basis of EFdA resistance and it is not known whether viruses with reduced sensitivity to EFdA would be cross-resistant to the NRTIs currently used in therapy.

In this study, we explored the influence of pre-existing NRTI resistance-associated mutations, and identified several novel mutations, associated with development of resistance to EFdA. We selected viruses resistant to EFdA by serial passage of HIV-1, initiating passages with WT virus, or with virus resistant to TFV, 3TC/FTC or AZT. We found that M184V was included in all selected EFdA-resistant strains. Mutations also appeared in the RT connection domain (R358K and E399K) and the RNase H domain (A502V), although these changes did not appear to compensate for loss of fitness nor significantly affect resistance when alone. We were able to identify a double mutant virus with a moderate-level of resistance, A114S/M184V (~25-fold).

## RESULTS

### Virus breakthrough during serial passage of viruses in increasing EFdA concentrations

Serial passages were initiated by infecting MT-2 cells with WT (xxLAI), K65R, M184V or D67N/K70R/T215F/K219Q stock virus in the presence of EFdA and followed as described in Materials and Methods. All EFdA-selected viruses, regardless of the initial strain or number of passages, induced ≥ 75% syncytia formation in untreated MT-2 cells within 7 days of infection.

The ability of the viruses to grow in the presence of EFdA was assessed by time to viral breakthrough (defined as 75% syncytia formation) at each passage. This time to breakthrough and the concentration of EFdA in each passage is shown in Figure 1. For WT, M184V, and D67N/K70R/T215F/K219Q, virus breakthrough typically occurred after approximately 7 days. As EFdA concentrations were raised in passages 8 and 9, time to virus breakthrough increased, until virus replication was no longer observed in passage 10. Passage 10 corresponded to an EFdA concentration of 550 – 800 nM (Figure 1A, C and D).

**FIG. 1.**
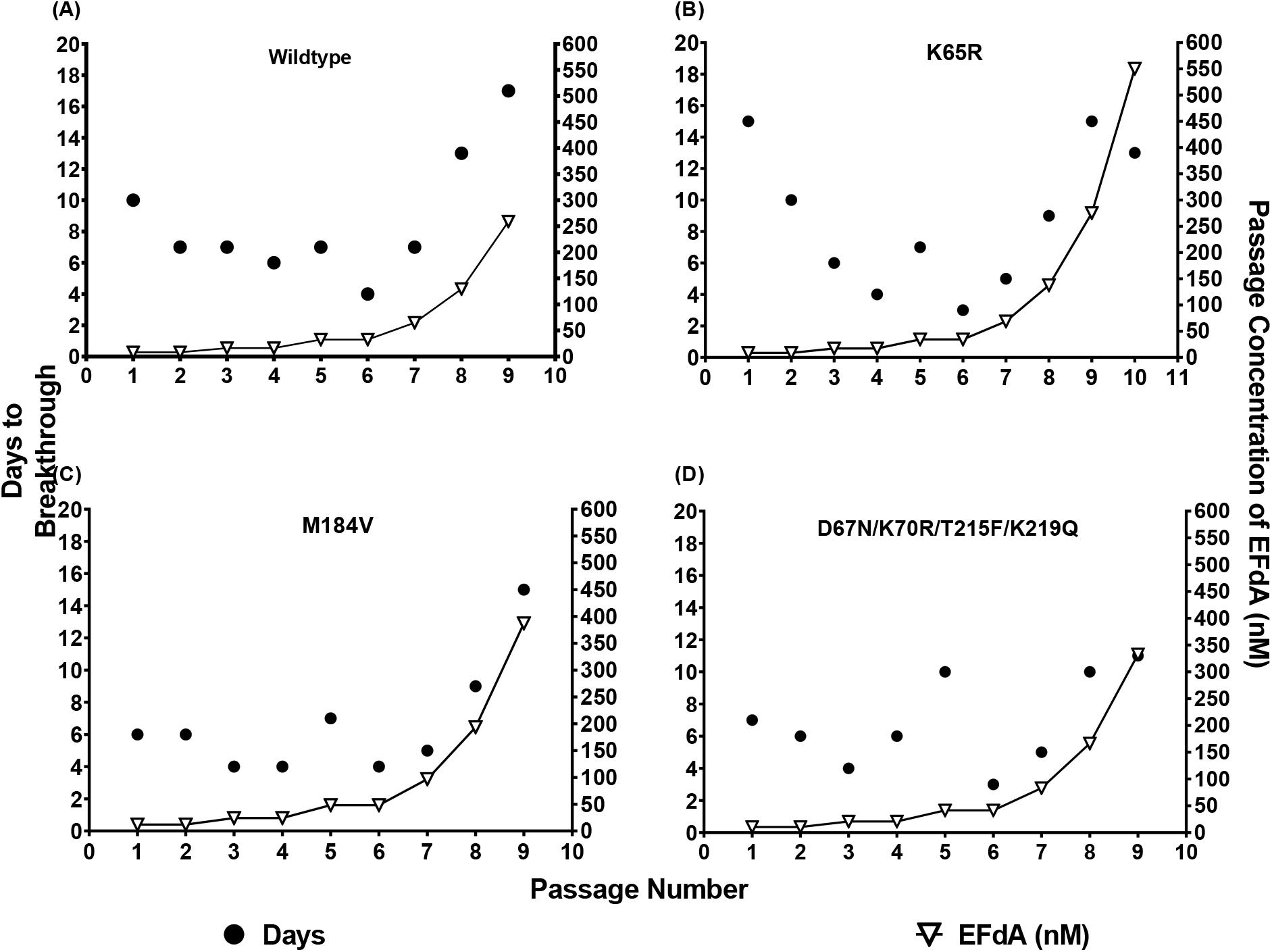
Selection of resistance to EFdA by serial passage. MT-2 cells were infected with 200 TCID50 of HIV-1 in media supplemented with EFdA. Cells were split 1:10 every 2 – 3 days and the supernatant replaced with fresh media supplemented with EFdA. Time to breakthrough was determined as the number of days required for ≥ 75% syncytia formation, at which point supernatants were harvested, assayed for p24 content and used to infect the subsequent passage. EFdA concentration was doubled every second passage until P_6_, and every passage thereafter. Results represent a single trial for each passage. Passages were initiated with (A) WT (xxLAI), (B) K65R, (C) M184V, and (D) D67N/K70R/T215F/K219Q.

The behavior of K65R deviated from the other mutants studied in two ways. Firstly, there was a reduction in the time to breakthrough from passages 1 – 4, likely associated with the loss of K65R that was lost as early as P_2_ (data not shown) and was not detected at passage 6 (Table 1, Figure 1), which confers hypersensitivity to EFdA (Michailidis et al., 2013). The time to breakthrough then increased as EFdA concentrations were raised, as seen with other viruses. These passages eventually terminated at passage 16 at a final concentration of ~35,000 nM EFdA (Figure 1B).

**Table 1.** Amino acid mutations in WT-, K65R-, M184V- and D67N/K70R/T215F/K219Q-derived viruses during serial passage in progressively increasing concentrations of EFdA

| Virus | Passage | Sequencing<sup>a</sup> | Proportion of Sequence Population (%) |
| --- | --- | --- | --- |
|  |  | Amino Acid Mutations<sup>b</sup> |  |
| WT (xxLAI) | 0<sup>c</sup> | None | 100 |
|  | 6 | M184I | 100 |
|  | 9 | M184I | 36.4 |
|  |  | M184I, E399K | 31.8 |
|  |  | M184V | 18.2 |
|  |  | None | 13.6 |
| K65R | 0<sup>c</sup> | None | 100 |
|  | 6 | R65K, M184I | 90.9 |
|  |  | R65K, M184V | 9.1 |
|  | 9 | R65K, M184V, A502V | 43.5 |
|  |  | R65K, M184I | 34.8 |
|  |  | R65K, M184V | 17.4 |
|  |  | R65K, M184I, A502V | 4.3 |
|  | 10 | R65K, A114S, M184V, A502V | 90.9 |
|  |  | R65K, A114S, M184V | 4.5 |
|  |  | R65K, A114S, A502V | 4.5 |
| M184V | 0<sup>c</sup> | None | 100 |
|  | 6 | R358K | 45 |
|  |  | None | 55 |
|  | 9 | R358K | 90.5 |
|  |  | None | 9.5 |
|  | 10 | R358K | 72.7 |
|  |  | R358K/A502V | 27.3 |
| D67N/<br>K70R/<br>T215F/<br>K219Q | 0<sup>c</sup> | None | 100 |
|  | 6 | M184V | 66.7 |
|  |  | None | 33.3 |
|  | 9 | M184V | 18.2 |
|  |  | M184V, E399K | 81.8 |

<sup>a</sup>Population and clonal sequencing of viral RNA from passage supernatants was performed as described in Materials and Methods.
<sup>b</sup>Relative to unpassaged stock virus of the appropriate strain
<sup>c</sup>Unpassaged stock virus

In every case, the passaged viruses were able to replicate in greater EFdA concentrations than the parental virus, indicating that some degree of EFdA resistance had developed. The most significant resistance appeared in the passages initiated with K65R virus.

### Infectivity of EFdA-passaged virus supernatants

To determine whether the passaged viruses had undergone significant changes in infectivity, independent of their potential resistance to EFdA, the infectivity of passaged isolates was compared to the infectivity of the unpassaged starting virus (P_0_), using the P4R5 MAGI reporter cell line in the absence of EFdA. For WT-, K65R- and M184V-derived viruses, the passaged strains demonstrated higher infectivity than the unpassaged P_0_ (Figure 2A - C). The D67N/K70R/T215F/K219Q viruses showed similar replication efficiency to the parental virus (Figure 2D). These increases in viral infectivity suggest that an increase in overall viral fitness may contribute to enhanced replication in the presence of EFdA for WT-, K65R- and M184V-derived viruses.

**FIGURE 2.**
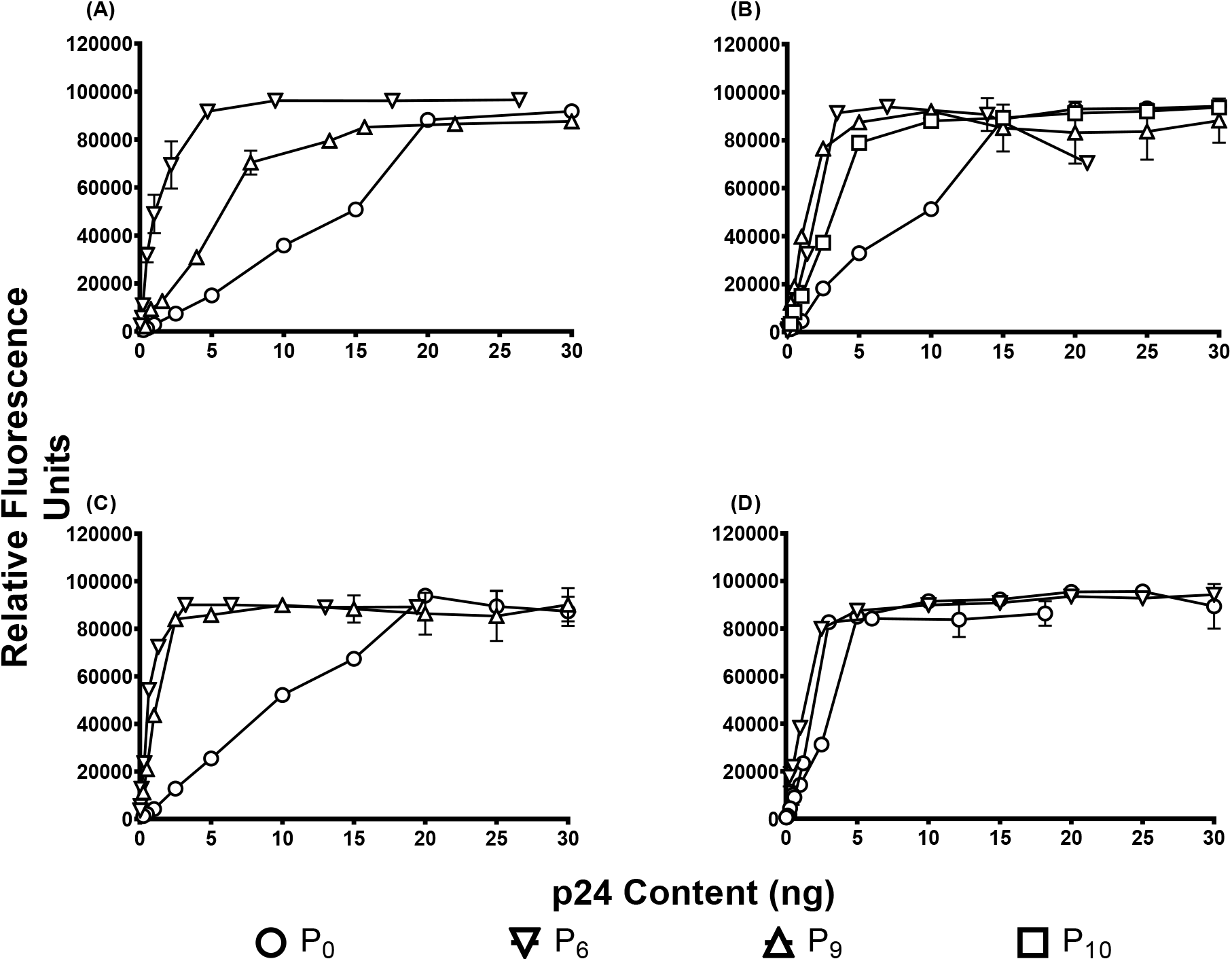
Infectivity of viruses selected during serial passage in EFdA. Untreated P4-R5 MAGI cells were infected with virus supernatants from P_0_, P_6_, P_9_ or P_10_ (where applicable). After 48 h, cells were lysed and β-galactosidase activity assessed. Virus infection is shown as relative fluorescence units. Results are the mean and S.D. of one experiment with three replicates. (A) WT (xxLAI), (B) K65R, (C) M184V, and (D) D67N/K70R/T215F/K219Q.

### Dose response of EFdA-passaged viruses to EFdA

To evaluate the degree of resistance selected in the passaging experiments, sensitivity to EFdA was determined for virus supernatants from P_6_ onwards. Data are presented for the viruses obtained from P_6_, P_9_ and, in one case, P_10_. P_6_ was the first passage for which extensive sequencing analysis was performed, P_9_ was the effective endpoint for the WT-, M184V- and D67N/K70R/T215F/K219Q-derived viruses, and P_10_ was chosen as the endpoint for the K65R-derived strain. For each of the viruses studied, the later passage viruses demonstrated increased EC_50_ values compared to WT and earlier passage viruses; however, the final EC_50_ increased compared to starting, as expected (Figure 3 A-D).

**FIGURE 3.**
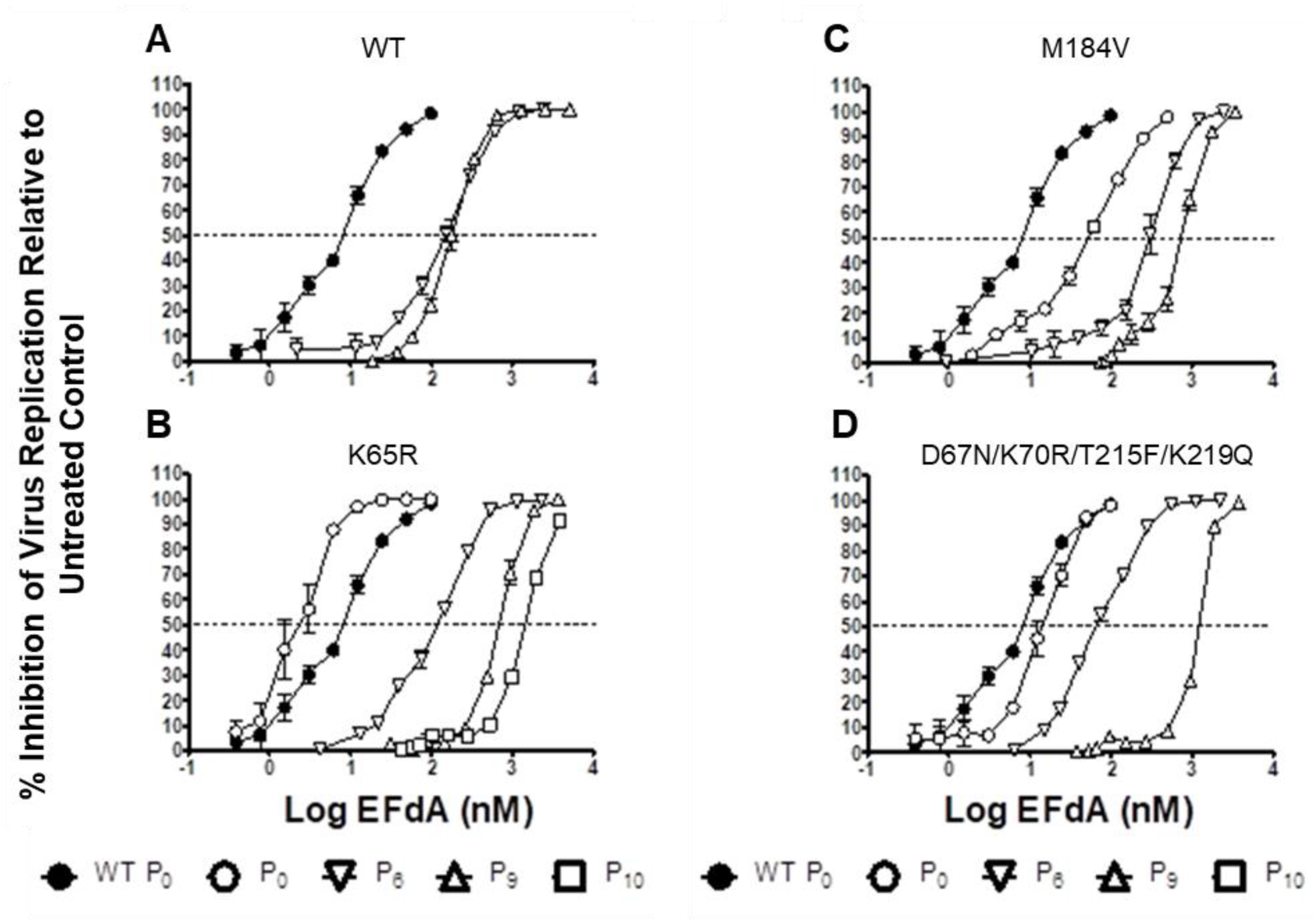
EFdA dose response for viruses selected during serial passage in EFdA. P4-R5 MAGI cells were infected with virus supernatants from P_0_, P_6_, P_9_ or P_10_ (where applicable) and treated with EFdA. Following incubation for 48 h, cells were lysed and β-galactosidase activity assessed. (A, B, C, D) Inhibition of virus replication relative to infected untreated cells. Dashed line represents 50% inhibition of viral replication. Results are the mean and S.D. of one experiment with three replicates.

### Development of amino acid mutations in reverse transcriptase during serial passage of viruses with EFdA

Clonal sequencing confirmed the identity of the P_0_ stock viruses. Similar sequencing was carried out on clones from various passages to identify any changes in RT that may have arisen during replication in the presence of EFdA. Table 1 summarizes data for P_6_ and P_9_ isolates for all parental viruses, and viruses from P_6_, P_9_, and P_10_ for K65R.

The WT P_6_ population sequencing revealed only M184I (Table 1); by P_9_, approximately 30% of sequences were M184I, 30% were M184I/E399K, and the remainder was divided between M184V and WT. Consistent with the initial passage data, sequencing data showed that all clones of the K65R P_6_ virus reverted the K65R mutation back to WT. The reversion occurred rapidly, as K65R was not seen in the P_2_ population. Over 90% of clones additionally contained M184I, with the remainder M184V. Sequencing data also showed that the proportion of M184V increased to ~one-quarter of the population by P_8_. The virus harvested following P_9_ had a more heterogeneous population, with most sequences containing M184V/A502V or M184I. By P_10_, the diversity decreased dramatically, and A114S/M184V/A502V became the dominant sequence. A114S/M184V and A114S/A502V were found at much lower frequency. The A114S/M184V/A502V mutation remained dominant through subsequent passages, reaching 100% of sequences in P_12_ and P_13_.

The only novel mutation gained during passages that started with M184V was R358K, which became increasingly dominant as passaging progressed. Sequencing of the virus that broke through P_10_ revealed that two-thirds of the sequences had R358K alone, while 27.3% of the population contained a combination of R358K and A502V in addition to the present M184V mutation.

Passage of the D67N/K70R/T215F/K219Q virus led to the rapid emergence of M184V (dominant in passage 6). This was joined by E399K (M184V/E399K was dominant in passage 9) in late passages. None of the starting mutations were lost during passage.

These passaging and sequencing experiments revealed a variety of mutations associated with EFdA resistance. Independent of the starting sequence, resistance was associated with mutations at residue M184, and the highest levels of resistance required an additional A114S mutation.

### Sensitivity of molecular clone viruses to EFdA

To confirm that the mutations identified by sequencing were capable of conferring resistance to EFdA, molecular clones representing the major mutant genotypes that developed during passage in EFdA were produced in a WT (NL4-3) backbone and characterized for their infectivity relative to WT. Individual mutants A114S, R358K, E399K, and A502V had no effect on resistance to EFdA. The only resistant individual mutant was M184V, which conferred about 8-fold resistance to EFdA, consistent with previous reports (Kawamoto et al., 2008b; Oliveira et al., 2017) (Figure 4). Addition of the A114S further increased the EFdA resistance, as seen in the cases of A114S/M184V (24-fold), A114S/M184V/R358K (28-fold), and A114S/M184V/A502V (25-fold). The R358K or A502V mutations by themselves had almost no effect on EFdA resistance (Figure 4). These data confirm that several of the mutations identified in the passaged viruses do confer resistance to EFdA, with the highest levels of resistance found in the concomitant presence of the A114S and M184V mutations.

**FIGURE 4.**
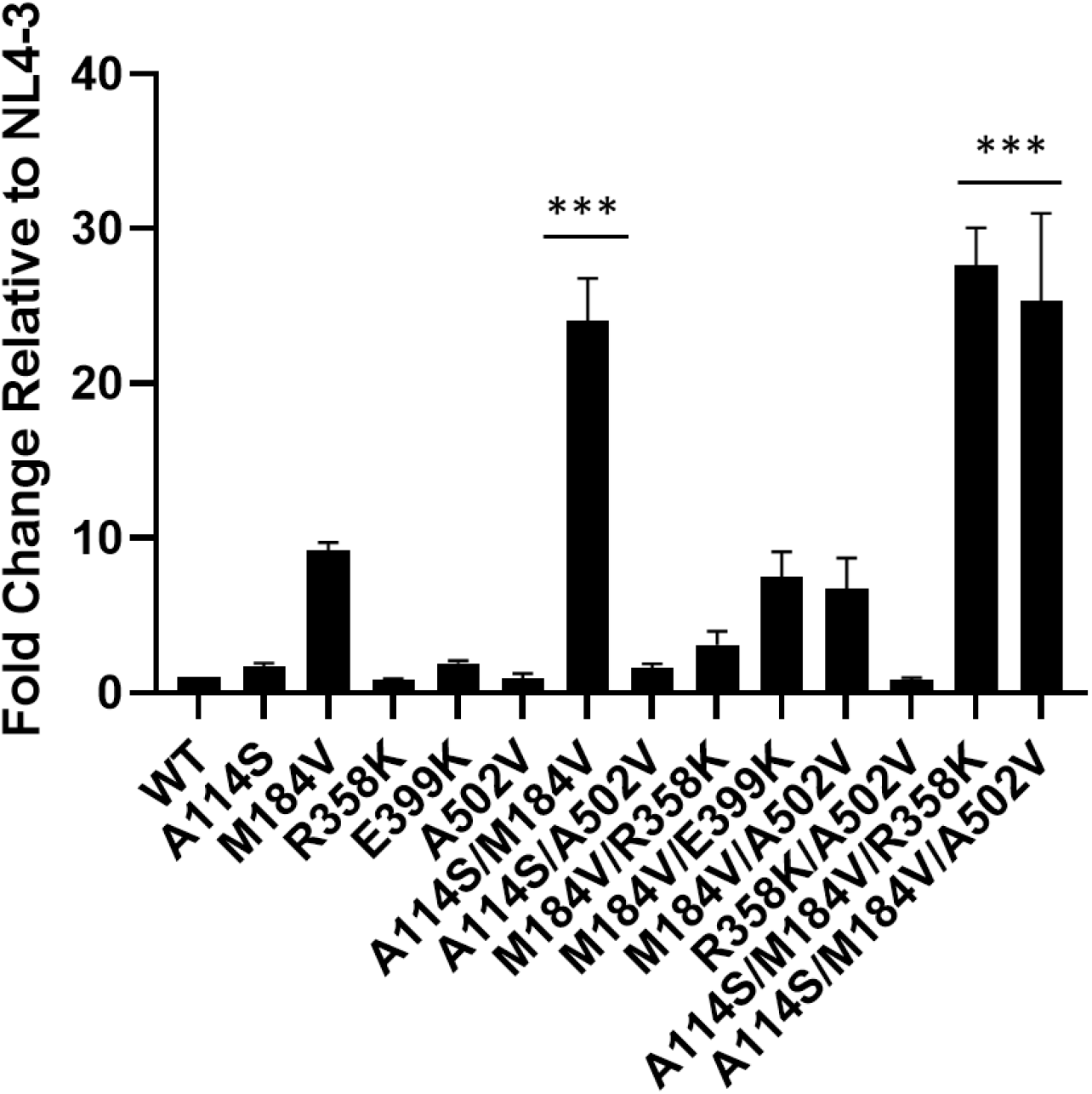
Fold-change EC_50_s relative to NL4-3. TZM-GFP cells were pre-treated with EFdA and infected after 24h. GFP positive cells (infected cells) were counted in varying concentrations of EFdA. Mutants were normalized to WT infection to produce fold change. EFdA dose response curves were produced for each mutant and the EC_50_s calculated. ***=p<.0001 Statistical significance was determined using a one-way ANOVA with Tukey’s post-test. Results are the mean and S.D. of four experiments with duplicates.

### Replication characteristics of molecular clone viruses

To determine if the mutations had an impact on viral fitness, viruses were tested in single cycle replication assays. There were no statistically significant differences in the fitness of connection domain mutants R358K and E399K or RNase H mutant A502V. However, there was a decrease in fitness of the A114S-containing mutants, A114S/M184V and A114S/M184V/A502V, as compared to WT (Figure 5). This is consistent with the biochemical data below, where a decrease in specific activity was observed for the RTs with the corresponding mutations.

**FIGURE 5.**
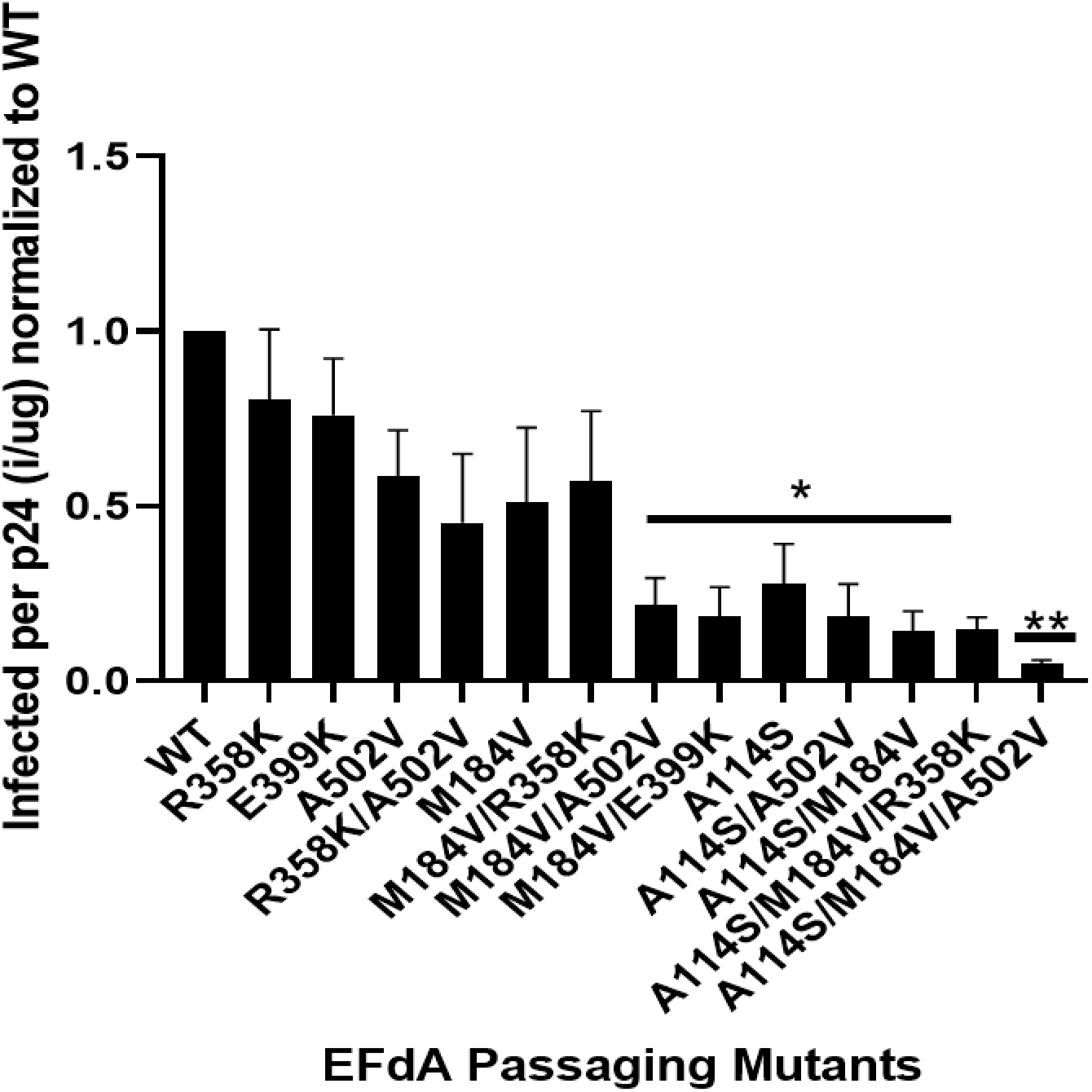
Single-round replication assays using TZM-GFP cells infected with individual mutants. An ELISA was also performed on the virus to determine amount of p24 (total virus in medium). The ratio of infected cells per p24 was then calculated and normalized to WT. Statistical significance was determined using a one-way ANOVA with Dunnet’s multiple comparison test (*=p<.05, **=p<.01). Results are the mean and S.D. of four experiments in triplicates.

### Steady State Kinetics and EFdA susceptibility of Mutant Reverse Transcriptases

In order to further understand the effect of the mutations on the DNA polymerase activity of RT, we performed steady state kinetics to determine the catalytic efficiency ratio, k_cat_/K_m_, for various RT mutants. We cloned, expressed, and purified the mutant RTs listed in Table 2. We found that in the presence of the A114S mutation there was a consistent decrease in the catalytic efficiency ratio k_cat_/K_m_ compared to WT RT: a 2.1 decrease for A114S and a 6.5 for A114S/M184V/A502V. Of note, the k_cat_/K_m_ for the A502V and M184V single mutants was comparable to that of the WT enzyme (Table 2).

**Table 2.** Steady-State Kinetics for Reverse Transcriptase Mutants

|  | WT RT | A114S | M184V | R358K | E399K | A502V | 184V/358K/502V | 114S/184V/502V |
| --- | --- | --- | --- | --- | --- | --- | --- | --- |
| K<sub>m</sub> (nM) | 1448 | 2758 | 1791 | 2008 | 1795 | 1110 | 2327 | 6659 |
| Fold Change K<sub>m</sub> | 1 | 1.9 | 1.2 | 1.4 | 1.2 | 0.8 | 1.6 | 4.6 |
| Fold Change k<sub>cat</sub>/K<sub>m</sub> | 1 | 2.1 | 1.2 | 1.4 | 1.2 | 1 | 1.3 | 6.5 |

## DISCUSSION

NRTIs are the most widely used therapeutics to treat HIV infection. As such, NRTI-resistant HIV variants are becoming increasingly prevalent in the HIV-1-infected population. EFdA has potential as salvage therapy for patients infected with NRTI-resistant mutants of HIV and as first line therapy for naïve HIV-infected individuals, owing to its potential suitability for long interval dosing. As such, we were interested in identifying and characterizing EFdA-resistant mutations that might arise during exposure of WT HIV, and especially NRTI-resistant virus variants, to EFdA.

Regardless the type of starting genotype virus, M184I/V ended up consistently to be the predominant mutation that arose during passages through EFdA. The M184V mutation confers high-level resistance to both 3TC and FTC (Petrella et al., 2004; Schinazi et al., 1993; Tisdale et al., 1993) (Sarafianos et al., 1998), but only low-level resistance to EFdA (Kawamoto et al., 2008a; Maeda et al., 2014). Our present data confirm the latter, with M184V conferring only 8-fold resistance to EFdA. Interestingly, M184V is also the primary resistance mutation selected during serial passage of WT HIV with related compounds EdA and Ed4T suggesting that M184 is critical to the activity of 4’-ethynyl modified nucleoside analogues (Kawamoto et al., 2008a; Nitanda et al., 2005).

Due to the widespread use of 3TC and FTC in HIV treatment, the M184V mutation is already present in many treatment-experienced patients. Since this mutation confers relatively low-level resistance to EFdA, we were interested to determine whether M184V virus could develop increased resistance during replication in the presence of EFdA.

The D67N/K70R/T215F/K219Q mutant, is highly resistant to AZT (Kellam et al., 1992; Larder and Kemp, 1989). Similar to another AZT-resistant mutant, M41L/T215Y, that was previously shown to display marginal resistance to EFdA (Kawamoto et al., 2008a), we found that D67N/K70R/T215F/K219Q had a 1.8-fold increase in EFdA resistance compared to WT. As we have previously shown, the excision unblocking mechanism of resistance is not a major challenge for EFdA: although EFdA can indeed be excised, the efficiency of reincorporation is so high, that the net result is no significant overall excision (Michailidis et al., 2014b). The K65R mutation confers resistance to TFV and is the mutation responsible for virological failure in TFV-based therapies. K65R is also cross-resistant or selected during therapy with ABC, ddI, and 3TC/FTC (Bazmi et al., 2000; Brenner and Coutsinos, 2009; Eggleton and Nagalli, 2020; Miller, 2004; Naeger and Struble, 2006; Zhang et al., 1994). We previously showed that the K65R variant is hypersensitive to EFdA, with up to five-fold lower EC_50_ compared to WT (Kawamoto et al., 2008a; Maeda et al., 2014; Michailidis et al., 2013). Our current results are consistent with this finding, as the K65R mutation was rapidly lost during passage in EFdA. The rapid reversion of this mutation leads not only to a reduction in sensitivity to EFdA, but also confers increased replication fitness to the virus. Collectively, the K65R, M184V, and D67N/K70R/T215F/K219Q, possess either increased sensitivity to EFdA, or slight (less than 10-fold) increased resistance.

Our results demonstrate the difficulty of selecting for resistance to EFdA. Even when starting the passages using different starting genotypic backgrounds, there was consistent appearance of the M184V mutation that by itself gave modest resistance 8-fold. Here we have identified a novel EFdA resistance mutation, A114S, that when added to M184V enhances EFdA resistance to about 25-fold. Importantly, this mutation, especially in the background of M184V, appears to have a negative impact on viral fitness.

This was confirmed using both viral fitness data as well as biochemical data with purified enzymes (Figure 5 and table 2). Specifically, it seems that the decrease in fitness is likely due to decreased binding of incoming dNTPs, as judged in all cases where A114S mutation was present. As both 184 and 114 residues are located at opposing sides of the EFdA binding pocket, we speculate that the bulkier mutant residues (V184 and S114) impinge into the substrate envelope of the 4’-ethynyl pocket, thus causing decrease in dNTP binding (increase in k_m_) and decrease in viral fitness.

In general, it appears that the development of EFdA resistance begins with mutations in the binding site; in our experiments, as well as in a previous report (Maeda et al., 2014), the initial mutation was M184I. We found that M184I was replaced at later passages by M184V, a mutation with superior replication capacity (Frost et al., 2000; Keulen et al., 1997; Schuurman et al., 1995). RT connection domain mutations (R358K and E399K) were selected at later passages. These residues are proximal to G359-A360 and K395-E396, respectively, in the RNAse H primer grip region that interacts with the DNA primer strand (Julias et al., 2003; Sarafianos et al., 2001). While R358 may form a weak hydrogen bond with the phosphate backbone of the primer strand (Ding et al., 1998), any effect of R358K or E399K will only be minor, and likely through minor structural adjustments affecting the position of nearby residues. Studies have demonstrated that R358K is selected both in NRTI-treated patients and during passage with NRTIs in tissue culture (Brehm et al., 2007; Delviks-Frankenberry et al., 2008; Lengruber et al., 2011; Tachedjian et al., 1998; von Wyl et al., 2010). However, any contribution of R358K to NRTI resistance is minimal and clinically it is likely selected as a pre-existing polymorphism, as it is present in 7.1% of treatment-naïve patients (Rhee, 2003; von Wyl et al., 2010). E399 is located within a cluster of several highly conserved tryptophan residues that are involved in RT dimerization (Chiang et al., 2012; Tachedjian et al., 2003), so E399K may have very small effects on stabilizing the heterodimer RT. We found no significant variation in EFdA resistance between M184V/E399K or M184V/R358K and M184V alone. These results indicate that neither R358K nor E399K contribute directly to EFdA resistance. We initially considered that these mutations might provide a fitness benefit to the mutants when added to the M184V background, potentially compensating for fitness costs associated with the acquisition of M184V (Wainburg, 2004; Xu et al., 2011). However, we did not observe improved fitness in our single round replication assays.

A further late-appearing mutation was A502V, which emerged during passaging of K65R. Unlike R358 and E399 that are located in the connection subdomain of RT, A502 is located in the RNase H domain, near residues Y501 and I505, both of which interact with the DNA primer strand as part of the RNAse H primer grip region (Sarafianos et al., 2001). Again, A502V does not appear to contribute to EFdA resistance, as the M184V/A502V clone did not have significantly lower sensitivity than M184V/E399K or M184V/R358K. The M184V/A502V mutant subsequently acquired an A114S mutation, which coincided with a significant increase in EFdA resistance. While the A114S mutation has never been reported in the context of EFdA resistance, it is associated with resistance to both foscarnet and AZT *in vitro*, but may increase sensitivity to AZT *in vivo* by affecting the enzyme processivity and pyrophosphorolysis rates (Arion et al., 2000; Larder et al., 2006, 1987).

A114 is part of a group of residues in the dNTP/EFdA-TP binding site, including A113, Y115, Q151, and M184, which interact with the 3’-OH of incoming dNTPs (Cases-Gonzalez and Menéndez-Arias, 2005; Harris et al., 1998; Van Cor-Hosmer et al., 2012). While the main chain amide of A114 interacts with the dNTP triphosphate, the alanine side chain protrudes towards the 3’-OH of the sugar moiety (Cases-Gonzalez and Menéndez-Arias, 2005; Huang et al., 1998; Menéndez-Arias, 2008). The A114S mutation has a longer side chain, which extends closer to the 3’-OH of an incoming dNTP and reduces the available space in the binding pocket and can thus increase selectivity for the correct dNTP. Consistent with our data, the A114S mutation was also previously observed to decrease the DNA polymerase activity of RT (Cases-Gonzalez and Menéndez-Arias, 2005; Van Cor-Hosmer et al., 2012). Furthermore, the side chain of A114 forms part of a hydrophobic pocket, along with Y115, M184, F160, and D185, which interacts with the 4’-ethynyl group of molecules such as EFdA (Salie et al., 2016) and 4’Ed4T-TP (Michailidis et al., 2009; Yang et al., 2008). M184V may alter the shape of this pocket, causing steric hindrance with the 4’-ethynyl and reducing the affinity of RT for molecules with this functional group (Yang et al., 2008). A114S may also alter the shape at the opposite side of the hydrophobic pocket. By itself, the A114S molecular clone demonstrated a modest increase in EFdA resistance relative to WT. Higher resistance (~24-fold) was observed when A114S and M184V appeared together. This could be due to both synergistically interfering with the hydrophobic pocket, thus disrupting the 4’-ethynyl from stabilizing in the hydrophobic pocket and greatly reducing the enzyme affinity for EFdA. It is also possible that the A114S mutation contributes to a reversal of the translocation impairment imposed by EFdA after its incorporation in the primer terminus.

Nevertheless, A114S-containing mutants may not be a major problem in patients; A114 mutations are rarely seen in clinical samples. Indeed, as of June 2012, only 21 sequences in the Stanford HIV Drug Resistance Database have mutations at this residue, only one of which is A114S (Rhee, 2003).

A major concern when introducing any new NRTI to clinical use is the potential for cross-resistance with current NRTIs. Our results confirm that EFdA drives the selection of M184I/V, producing virus with high-level cross-resistance to both 3TC and FTC. However, EFdA is a potent inhibitor, even against M184I/V mutants; it retains the ability to inhibit replication of these mutants at therapeutic doses. Conversely, M184I/V increases AZT sensitivity in both WT and AZT-resistant backgrounds (Boucher et al., 1993; Boyer et al., 2002; Larder et al., 1995; Tisdale et al., 1993).

The selection and characterization of EFdA resistance mutations through viral passage revealed the difficulty in identifying mutations that confer high-level resistance to EFdA. Although mutations at M184 were selected, they conferred less than 10-fold resistance. In our experiments, the development of greater resistance to EFdA required the combination of multiple mutations, including both M184V and A114S. These changes allowed for enhanced resistance to EFdA.

In summary, the data presented here demonstrate that a significant barrier exists to HIV developing high-level resistance to EFdA. In addition, viruses that do develop high-level resistance to EFdA become highly sensitized to other NRTIs. These results demonstrate that EFdA has potential to be a highly effective therapeutic.

## MATERIALS AND METHODS

### Reagents

EFdA was synthesized by Life Chemicals (Burlington ON, Canada). Stock solutions (10 mM) of EFdA was prepared in dimethyl sulfoxide (DMSO) and stored in aliquots at −20 ^°^C. MT-2 cells (Boufford and Spongberg, 1983; Charneau et al., 1994; Haertle et al., 1988) were cultured in RPMI 1640 medium (Mediatech Inc, Manassas, VA), supplemented with 10% fetal bovine serum (FBS) (HyClone, Logan, UT), 2 mM L-glutamine (Mediatech Inc) and 100 U/ml penicillin, 100 μg/ml streptomycin (Mediatech Inc). P4-R5 MAGI cells were cultured in DMEM (Mediatech Inc), supplemented with 10% FBS, 2 mM L-glutamine, 100 U/ml penicillin, 100 μg/ml streptomycin and 1 μg/mL puromycin. HEK-293 cells (Graham et al., 1977) and HEK-293/17 (Pear et al., 1993) were cultured in DMEM supplemented with 10% FBS, L-glutamine and penicillin/streptomycin. TZM-GFP cells (Derdeyn et al., 2000; Platt et al., 2009, 1998; Rosa et al., 2015; Takeuchi et al., 2008; Wei et al., 2002) were cultured in DMEM (Corning) supplemented with 10% Serum Plus, 2 mM L-glutamine (ThermoFisher, Waltham, MA), 100 U/ml penicillin/streptomycin (ThermoFisher). Jurkat LTR-GFP CCR5+ Cells (JLTRG-R5) (Kutsch et al., 2004; Ochsenbauer-Jambor et al., 2006) were cultured in RPMI 1640 medium (Cytiva Life Sciences, Marlborough, MA) supplemented with 10% FBS and 100 U/mL penicillin/streptomycin.

### Generation of virus stocks and molecular clones

Initial K65R, M184V, and D67N/K70R/T215F/K219Q viruses were generated by site-directed mutagenesis on an xxLAI HIV-1 backbone using the QuikChange XL Site-Directed Mutagenesis kit (Agilent Technologies Inc, Santa Clara, CA), according to the manufacturer’s protocols.

Subsequently, 6 x 10^5^ 293-T cells were transfected with 10 μg of viral DNA using the PrimeFectimine Mammalian Transfection Reagent (PrimGen, Oak Park, IL). After 72 h incubation, HEK-293/T cell supernatants were harvested, filtered and used to infect 1.8 x 106 MT-2 cells. Infected MT-2 cells were incubated at 37 oC (5% CO2), inspected daily and infectious virus harvested at ≥ 50% syncytia formation. Emory’s Cloning Core was used to make individual mutants in the backbone of NL4.3. These mutant viruses were made using HEK-293/17 cells that were transfected with 6 μg of viral DNA using Xtreme-GENE HP (Roche, Basel, Switzerland) Transfection reagent. After 48 h incubation HEK-293/17 cell supernatants were harvested, concentrated overnight with a Lenti-X concentrator (Clontech) according to the manufacturer’s protocol.

### Determination of TCID_50_ values and p24 content

TCID50 values were determined by infecting 5 x 10^4^ MT-2 cells per well, in 96-well flat-bottom plates, with four-fold serial dilutions of virus stock. Three replicates were performed for each virus. Infected plates were inspected daily for syncytia formation; every three days, half of the supernatant was replaced with fresh media. The assay was terminated when no additional syncytia formation was noted for two days. The TCID_50_ was then calculated using the Reed-Muench method (Reed and Muench, 1938). The p24 content of each virus stock was determined using the HIV-1 p24^CA^ Antigen Capture Assay kit (SAIC-Frederick, Frederick, MA).

### Serial passage for selection of resistant virus

MT-2 cells were suspended at 2.5 x 10^5^ cells per mL in 10 mL of media containing EFdA. Initial EFdA concentrations were chosen based on the EC_50_ of WT stock virus (8.6 nM), with a similar amount chosen for the K65R virus and slightly higher concentrations for the D67N/K70R/T215F/K219Q (10 nM) and M184V (12 nM) strains. Passages were initiated by immediately adding 200 TCID_50_ of the appropriate unpassaged (P_0_) virus stock to the cells and mixing gently. Untreated cultures were initiated by infecting 2.5 x 105 MT-2 cells per mL with 200 TCID_50_ of P_0_ virus stock in 10 mL drug-free media. All passages and untreated cultures were grown in T-25 tissue culture flasks. Every 2 – 3 days, cells were mixed and replaced with fresh media containing the appropriate concentration of EFdA. Cultures were visually inspected every 1 – 2 days for the presence of syncytia. At ≥ 75% syncytia formation, culture supernatants were harvested, concentrated using Amicon Ultra Ultracel – 100K centrifugal filters (Millipore, Carrigtwohill, Co. Cork, Ireland) and syringe-filtered through 0.22 μM filters (Millipore). The p24 content of the resulting first passage (P_1_) supernatant was determined as described above. This procedure was followed for all subsequent passages, with P_N_ initiated by infecting 2.5 x 10^6^ MT-2 cells with P_N-1_ virus supernatant in media containing the appropriate concentration of EFdA. Untreated cultures were also initiated by infecting 2.5 x10^6^ MT-2 cells, in drug-free media, with P_N-1_ virus. As infectivity differences were expected between the P_0_ virus and the output strains from each passage, P_2_ and all subsequent passages were initiated by infecting cells with a p24 amount of P_N-1_ virus equivalent to the P_0_ virus p24 content used to initiate P_1_. The concentration of EFdA was doubled every two passages up to P_6_, after which point the amount of drug was doubled every passage. Passages lasting more than 60 days without syncytia formation were terminated and repeated, along with the previous passage. If no syncytia were noted in a repeat passage after 30 days, the passage was terminated and no further attempts were performed.

### Dose response to individual mutants

All individual mutants were cloned into NL4.3 using Emory’s Cloning Core. After each virus was generated as stated above, TZM-GFP cells were plated at 1 x 10^3^ cells/well in a 96-well plate and with serial-diluted EFdA starting at 1 nM and the cells and EFdA were incubated for 24 h. After, the cells were infected with virus and 1 μg/mL final concentration of DEAE Dextran and incubated for 48. The GFP positive cells were then counted using Cytation 5 (Biotek, Winooski, VT) with Gen5.5 Software. EC_50_ curves were then determined using Prism 5 (GraphPad) software.

### Viral Replication Assays

JLTRG-R5 cells were plated at 5 x 10^6^ cells/well in a 12 well plate. The plasmid NL4.3 and various mutants were transfected into JLTRG-R5 cells with 1 μg of viral DNA using Xtreme-GENE HP (Roche) Transfection reagent. After 48 h incubation, cells were imaged using Cytation 5 to quantify GFP sum signal, cells were then split and media was replaced. Cells were subsequently imaged and split every 2-3 days until day 20. After day 20, genomic DNA was extracted using DNeasy Blood & Tissue Kit (Qiagen, Valencia, CA). Genomic DNA was PCR amplified using primers: 5’-gaagaaatgaatttgccagg-3’ and 5’-ctcatgttcttcttgggc-3’ and Phusion DNA Polymerase Master mix (New England Biolabs, Ipswich, MA). DNA was sent for Sanger Sequencing to check for reversion mutations.

### Specific Infectivity

TZM-GFP cells were plated at 10,000 cells/well in a 96-well plate and incubated for 24 h. After, cells were infected with the varying concentrations of the virus and a 1 μg/mL final concentration of DEAE-dextran and incubated for 48 h. The GFP positive cells were then counted as described above. The p24 content of each virus was also determined using an ELISA.

### Steady State Kinetics and *in vitro* IC_50_s

HIV-1 RT and mutants were expressed and purified as described previously (Bauman et al., 2008; Kirby et al., 2012; Michailidis et al., 2009; Ndongwe et al., 2012; Sarafianos et al., 2003; Schuckmann et al., 2010). RT was expressed in JM-109 cells (Invitrogen) and purified by nickel affinity chromatography and Mono Q anion exchange chromatography. Steady state kinetic parameters, k_m_, for incorporation of EFdA-MP were determined using plate-based assays measuring an 18 nucleotide primer annealed to 100 nucleotide DNA template (Kankanala et al., 2017; Singh et al., 2012; Tang et al., 2019, 2017; Vernekar et al., 2017; Wang et al., 2018). The reactions were carried out in RT buffer with 6 mM MgCl_2_, 40 nM Td100/Pd18, and 10 nM RT in a final volume of 20 μl for 30min at 37 °C, and arrested by 100 mM of EDTA. The QuantiFluor dsDNA System (Promega) was used to quantify the amount of formed double-stranded DNA. Reactions were read at ex/em 504/531 nm in a PerkinElmer EnSpire Multilabel plate reader. k_m_ were determined graphically using the Michaelis-Menten equation using Prism Software.

### Sequencing of passaged viruses

Viral RNA was purified from supernatants using the QIAamp Viral RNA Mini kit (Qiagen, Valencia, CA), the concentration determined with a Spectronic BioMate*3 UV spectrophotometer (Thermo Scientific, Waltham, MA) and 500 ng used as the template for cDNA synthesis. First-strand PCR was performed using random hexamer primers and the SuperScript III First-Strand Synthesis System for RT-PCR (Invitrogen, Carlsbad, CA). The resulting cDNA was PCR amplified using HIV-1 LAI-specific primers ABR-RT-OF (1763 5’-GGAGCCGATAGACAAGGAACTG-3’) and ABR-RT-OR2 (3863 5’-GGCTACTATTTCTTTTGCTACTACAGG-3’). These primers anneal to the 3’ end of gag and the 5’ end of integrase, respectively, and generate a 2127 bp product spanning the full length of the reverse transcriptase gene. PCR was performed using the Expand High Fidelity PCR System dNTPack (Roche Diagnostics GmbH, Mannheim, Germany), with 4.5 mM MgCl_2_ and 3.5 U of enzyme mix used for each reaction. Reactions were run in a PCR Sprint bench-top PCR cycler (Thermo Electron) with an initial denaturation of 3 minutes at 94 ^°^C, followed by 30 amplification cycles, each consisting of 30 seconds at 94 ^°^C, 45 seconds at 58 ^°^C and 150 seconds at 72 ^°^C. Following a seven-minute final extension at 72 ^°^C, samples were used immediately or stored at −20 ^°^C.

PCR products were separated by electrophoresis on a 1% agarose gel, and the 2127 bp band harvested using the QIAquick Gel Extraction kit (Qiagen, Valencia, CA). The concentration of DNA was determined by spectrophotometry, and 500 ng of sample was submitted to ACGT Inc (Wheeling, IL) for full-length, double-stranded population sequencing. The P_0_ consensus sequences from each virus were aligned in Clustal X-2 (Larkin et al., 2007) with an independent HIV-1 LAI nucleotide sequence (Accession NC_001802) to ensure the stock viruses did not contain unexpected mutations. Nucleotide consensus sequences from each passage were aligned to the appropriate P_0_ consensus. Chromatograms were also inspected visually using Chromatogram Explorer and DNABaser (Heracle BioSoft S.R.L., Pitesti, Romania) for the presence of heterogeneous peaks and minority sequence populations not detectable in the consensus sequence.

### Clonal sequencing of full-length reverse transcriptase gene PCR products

Approximately 105 ng of full-length PCR product was ligated into the pGEM-T Vector System (Promega, Madison, WI), at a 3:1 molar ratio of insert:vector and incubated overnight at 4 ^°^C. Ligations were transformed into MAX Efficiency DH5α competent cells (Invitrogen, Carlsbad, CA) by heat shock. Blue-white screening was used to select clones with successful ligations, and plasmids containing the full-length reverse transcriptase gene were isolated with the QIAprep Spin Miniprep kit (Qiagen, Valencia, CA). A minimum of 20 clones from each sample were sequenced. Primers ABR-RT-OF and ABR-RT-OR2 were used to sequence the 5’ and 3’ ends of reverse transcriptase, while an internal portion of the gene was sequenced with primer ABR-RT-IF (2211 5’-CAGAGATGGAAAAGGAAGGG-3’). Clones were aligned with the appropriate P_0_ stock virus consensus in Clustal X-2, and the proportion of sequences with the novel substitutions was determined.

## ACKNOWLEDGMENTS

We acknowledge the key experimental contribution by Aaron B Reeve in the viral passage experiments and also in the initial stages of writing a different version of the manuscript. As he is inaccessible to approve this version of the manuscript, he is currently not included in the author list, in compliance to the Journal’s editorial guidelines. The author list will be updated appropriately when he approves the submitted version. Michael Parniak was involved in the early research and writing processes; as he is deceased it has not been possible to obtain his approval of the final version of the manuscript.

This work was supported in whole or in part by NIH grants AI076119 to S.G.S. T32GM008367 (provided training funds for M.E.C.).

The following reagents were obtained through the AIDS Research and Reference Reagent Program, Division of AIDS, NIAID, NIH: MT-2 cells from Dr. Douglas Richman, P4-R5 MAGI cells from Dr. Nathaniel Landau; HEK-293 cells from Dr. Andrew Rice.

This study was supported in part by the Emory Integrated Genomics Core (EIGC), which is subsidized by the Emory University School of Medicine and is one of the Emory Integrated Core Facilities. Additional support was provided by the National Center for Advancing Translational Sciences of the National Institutes of Health under Award Number UL1TR000454. The content is solely the responsibility of the authors and does not necessarily reflect the official views of the National Institutes of Health.

